# Label-free imaging of collagen fibers in tissue slices using phase imaging with computational specificity

**DOI:** 10.1101/2021.11.19.469223

**Authors:** Masayoshi Sakakura, Virgilia Macias, Sohelia Borhani, André Kajdacsy-Balla, Gabriel Popescu

## Abstract

Evaluating the tissue collagen content in addition to the epithelial morphology has been proven to offer complementary information in histopathology, especially in oncology tumor staging and prediction of survival in cancer patients. One imaging modality widely used for this purpose is second harmonic generation microscopy (SHGM), which reports on the nonlinear susceptibility associated with the collagen fibers. Another method is polarization light microscopy (PLM) combined with picrosirius-red (PSR) tissue staining. However, SHGM requires expensive equipment and provides limited throughput, while PLM and PSR staining are not part of the routine surgical pathology workflow. Here, we utilize phase imaging with computational specificity (PICS) to computationally infer the collagen distribution of *unlabeled* tissue, with high specificity. PICS utilizes deep learning to translate quantitative phase images (QPI) into corresponding PSR images with high accuracy and inference speed of 200 milisecond per forward-pass through the model once trained. We developed a multimodal imaging instrument that yields both Spatial light Inference Microscopy (SLIM) and polarized light microscopy (PLM) images from the same field of view. Our results indicate that the distributions of collagen fiber orientation, length, and straightness reported by PICS closely match the ones from ground truth as defined by KL-divergence.

## Introduction

Collagen plays an important role in skin aging **[1]**, is used in surgical treatments of vitiligo **[1]**, and its degradation is known to be associated with proliferation of melanoma cells **[2]**. It has been shown that collagen fiber orientation can report on tumor aggressiveness via a tumor-associated collagen signature (TACS) **[3–6]**.

Collagen is the body’s most abundant protein and is used in a wide variety of biomedical applications ranging from bone reconstruction **[7]** to wound healing **[8]** and skin regeneration **[9]**. As a major component of the tumor microenvironment, collagen plays a critical role in cancer progression, angiogenesis, and metastasis, as well as resistance to radiation and chemotherapy **[10]**. The “tumor-associated collagen signatures” (TACS) model **[11]** describes various distribution patterns exhibited by collagen fibers in the vicinity of malignant cells, which can be used to infer tumor aggressiveness. Moreover, individual fiber properties including length, straightness, and orientation have all been shown to carry valuable information related to the progression of the disease **[12–14]**. Hence, end-to-end systems capable of quantifying collagen content and orientation in tissue sections can be of great utility for pathologists, oncologists, and cancer researchers.

Second harmonic generation microscopy (SHGM) maps the second-order nonlinear susceptibility *χ*^(2)^ associated with *non-centrosymmetric* molecules and, thus, probes fibrillar collagen structures with high specificity **[15]**. Collagen generates a strong second-harmonic signal compared to the surrounding cellular structures, resulting in high imaging contrast **[16–18]**. However, the low contrast of other cellular structures in SHGM images means that centrosymmetric structures, such as tumor boundaries are difficult to delineate and often other imaging modalities are required **[19]**. Importantly, due to the phase matching condition, imaging collagen via SHGM is limited to 2D. Recently, it has been shown that using phase-resolved data and a new physical model that takes into account the scattering and phase-matching condition, artifact-free *χ*^(2)^ 3D tomography can be obtained **[20]**.

Quantitative phase imaging (QPI) is a rising field in label-free optical imaging for biomedical applications **[21, 22]**. Unlike many stain-dependent imaging modalities, QPI solely relies on the intrinsic contrast from the density-dependent morphology of the sample, which has a direct correspondence to optical pathlength difference. As a result, QPI can detect transparent, weakly-scattering features from biological samples with nano-scale sensitivity **[23–30]**. QPI has been applied at a rapid pace in a variety of basic science **[31–39]** and clinical applications **[21, 40–46]**. Recently, it has been shown that QPI is extremely sensitive to collagen in histology slides. Comparing the QPI and SGHM data on the same fields of view, we found that QPI can extract collagen fiber orientation accurately and also provide the contextual structures, absent in SHGM **[5, 47]**. Spatial light interference microscopy (SLIM) is a highly sensitive QPI technique widely used for biomedical applications **[25, 48, 49]**. One of the appealing features of SLIM is that it can be implemented as a module attached to the camera port of a commercial phase-contrast microscope **[25, 38, 49]**. This versatility of SLIM installation allows multi-channel, correlative imaging, in combination with fluorescence **[50, 51]** or other modalities, such as polarization light microscopy, which is relevant to our study here. In addition, SLIM interfaces seamlessly with the motorized microscope and allows for whole slide QPI **[46]**. Since SLIM takes advantage of low-coherence light source into a completely common-path microscopy system, the acquired image from SLIM has a very low spatial and temporal noise level.

Recently, deep learning has shown tremendous potential in biophotonics in general and microscopy in particular **[52–56]**. Ozcan and colleagues have pioneered image to image translation approaches based on deep learning resulting in synthetic images of stains and fluorescence extracted from label-free data **[57–60]**. Our group has advanced a family of tools collectively referred to as phase imaging with computational specificity (PICS) which aims to achieve chemical specificity from unlabeled specimens using deep learning **[50, 51, 61–65]**. Thus, PICS allows for nondestructive imaging, of low exposure and chemical phototoxicity, while maintaining the specificity associated with exogenous contrast agents.

In this study, we extend the PICS framework to clinical relevance. We demonstrate that label-free analysis of collagen fibers can be performed with high specificity by combining SLIM with deep learning. We developed a multimodal imaging instrument that yields both SLIM and polarized light microscopy (PLM) images from the same field of view. Using conventional picrosirius-red (PSR) staining, which is specific to collagen in polarized microscopy, we acquired images of the collagen fibers and used them as ground truth **[66]**. On the same field of view, we acquired SLIM data, such that the resulting pairs of images were used for training the Efficient U-Net **[67]** to infer synthetic PSR images from QPI data. We validated the trained network over an unseen SLIM dataset to evaluate the network’s capability for label-free identification of collagen fibers. We further used a collagen fiber tracking algorithm, referred to as CT-FIRE **[19, 68, 69]** and showed that it can directly be applied on the PICS-generated collagen map without the need for PSR stains. Using CT-FIRE we further performed segmentation to detect individual fibers and analyze their statistical distribution in terms of length, straightness, and angle of orientation. Our results show a strong statistical similarity between the CT-FIRE output from the ground truth dataset and the corresponding distributions from the PICS inference.

The manuscript is structured as follows. 1) We present the SLIM/ PLM setup and the corresponding images acquired, 2) Efficient-U-Net architecture and the performance metrics, 3) CT-FIRE analysis of extracted collagen fibers, and 4) statistical validation of the collagen fiber distributions. In the Methods section, we elaborate on the optical setup and hardware used for SLIM and PLM, the PICS implementation, and the CT-FIRE algorithm for statistical validation.

## Results

### SLIM and PLM multimodal imaging system

We combined SLIM, which yields quantitative phase maps associated with the specimen **[25, 49]**, and polarization light microscopy (PLM), which reports on the birefringence associated with the PSR intensity distribution **[68, 70, 71]**. The configurations for both imaging modalities are discussed in more detail in the Methods section. Briefly, as shown in Figure 1, both SLIM and PLM modules are attached to the side ports of a commercial microscope. SLIM uses the phase contrast channel and consists of an external optical module that generates additional phase-shifts between the incident and scattered fields **[25, 49]**. PLM imaging is done under a cross-polarized brightfield optical path, in which only the regions with birefringence in the sample generate signal **[72]**.

**Figure 1:**
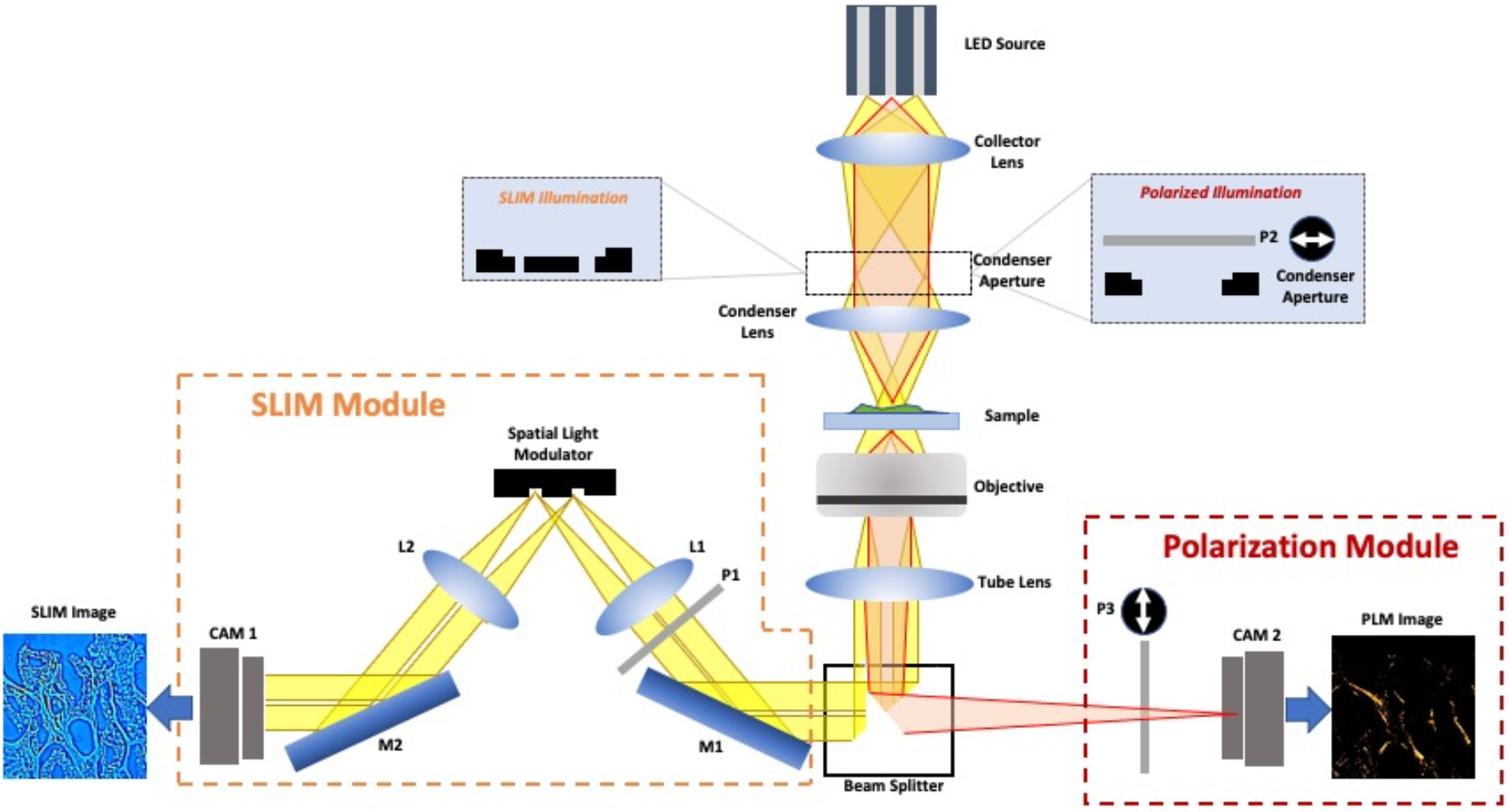
Schematic of dual-camera imaging setup involving SLIM and PLM. The dual-camera setup consists of a standard inverted optical microscope with two external modules attached on either side of the output port. The light-path can be switched by turning the beam-splitter. The condenser aperture for the SLIM is a simple dark-field illumination which is typical for phase-contrast microscopy. The condenser aperture used for polarized illumination consists of input polarizer P2 on top of a brightfield condenser aperture. The output linear polarizer P3 is placed in the polarization module such that its transmission axis is orthogonal to that of P2.

For PICS training and testing, we imaged a total of 217 cores from a prostate tissue microarray (TMA), resulting in a total of 1,000 fields of view imaged by both SLIM and PLM. Setting the overlap percentage between adjacent frames to 5 %, we digitally mosaicked individual fields of view to obtain images of the entire cores, each 900 μm diameter. **Figure 2** presents both the microscopic and the macroscopic images acquired under SLIM and PLM. The PLM channel allows us to localize and analyze the collagen fiber content with high specificity **[68, 70, 71]**, while SLIM reports on tissue morphology information without staining, providing contextual information, such as epithelium and non-collagen extracellular matrix **[50]**.

**Figure 2:**
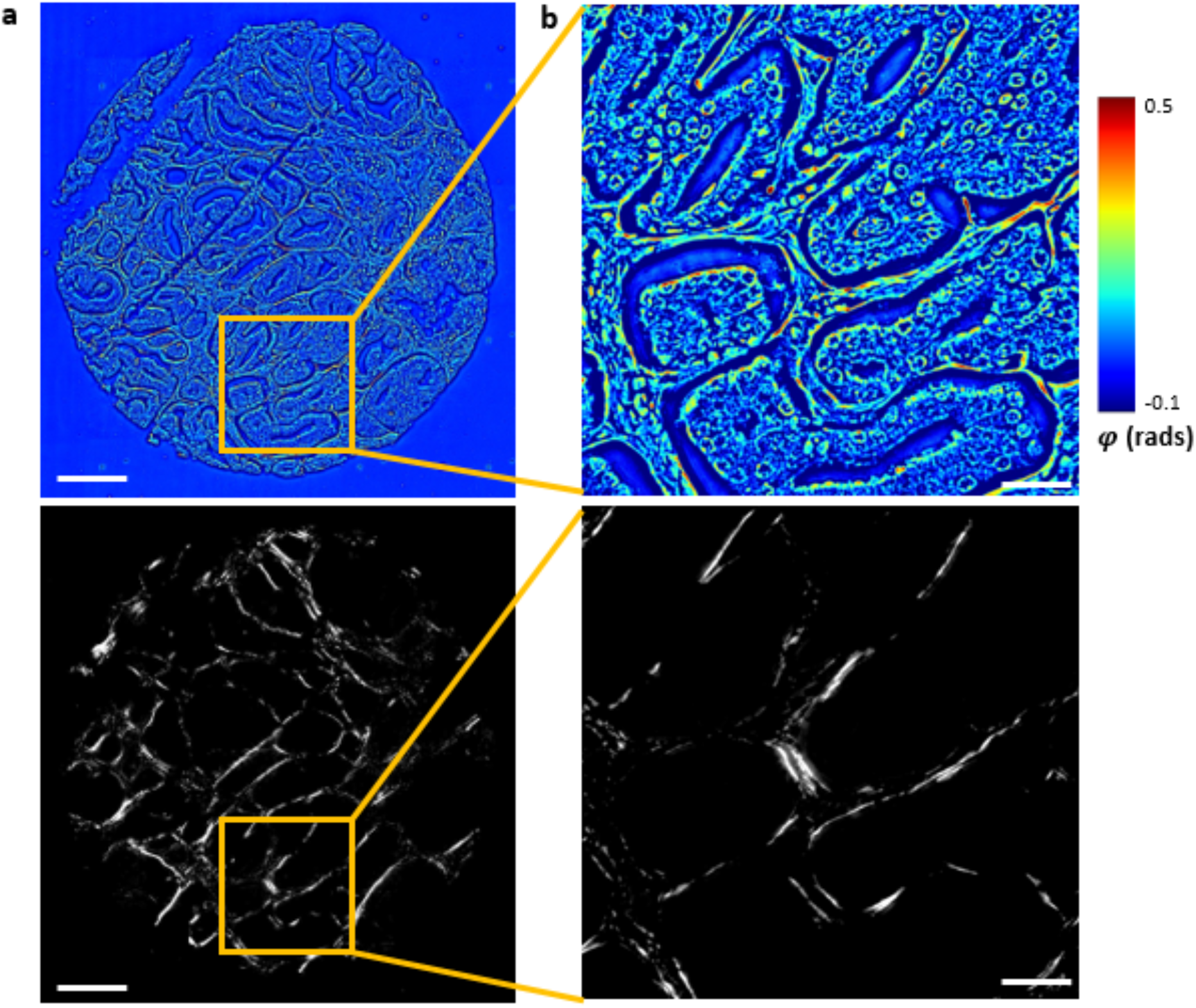
TMA core imaged with SLIM and PLM. **a.** A digitally stitched TMA core imaged with SLIM (top) and PLM (bottom). The white bar represents 200 microns in length. **b.** A single field of view from a core imaged under SLIM (top) and PLM (bottom). The white bar represents 50 microns in length.

### Procedures Prior to Network Training

The PICS method for collagen detection follows the workflow shown in **Figure 3a**. The experimental procedure first begins by acquiring the image dataset under both SLIM and PLM. We then discard the empty frames, i.e., the fields of view that contain only background and have no tissue content. An important part of the pre-processing is the proper registration of the images from the two channels. The MATLAB registration estimator app was initially used to infer a transformation matrix that geometrically transforms the images from the PLM camera to match those from the SLIM camera. The evaluated matrix was then used to transform all the collected PLM images to complete the registration process as shown in **Figure 3b**. The pre-processed image pairs are randomly assigned into either training, validation, or test set for E-U-Net training. Figure 3c presents the fiber extraction from a TMA core using the CT-FIRE.

**Figure 3:**
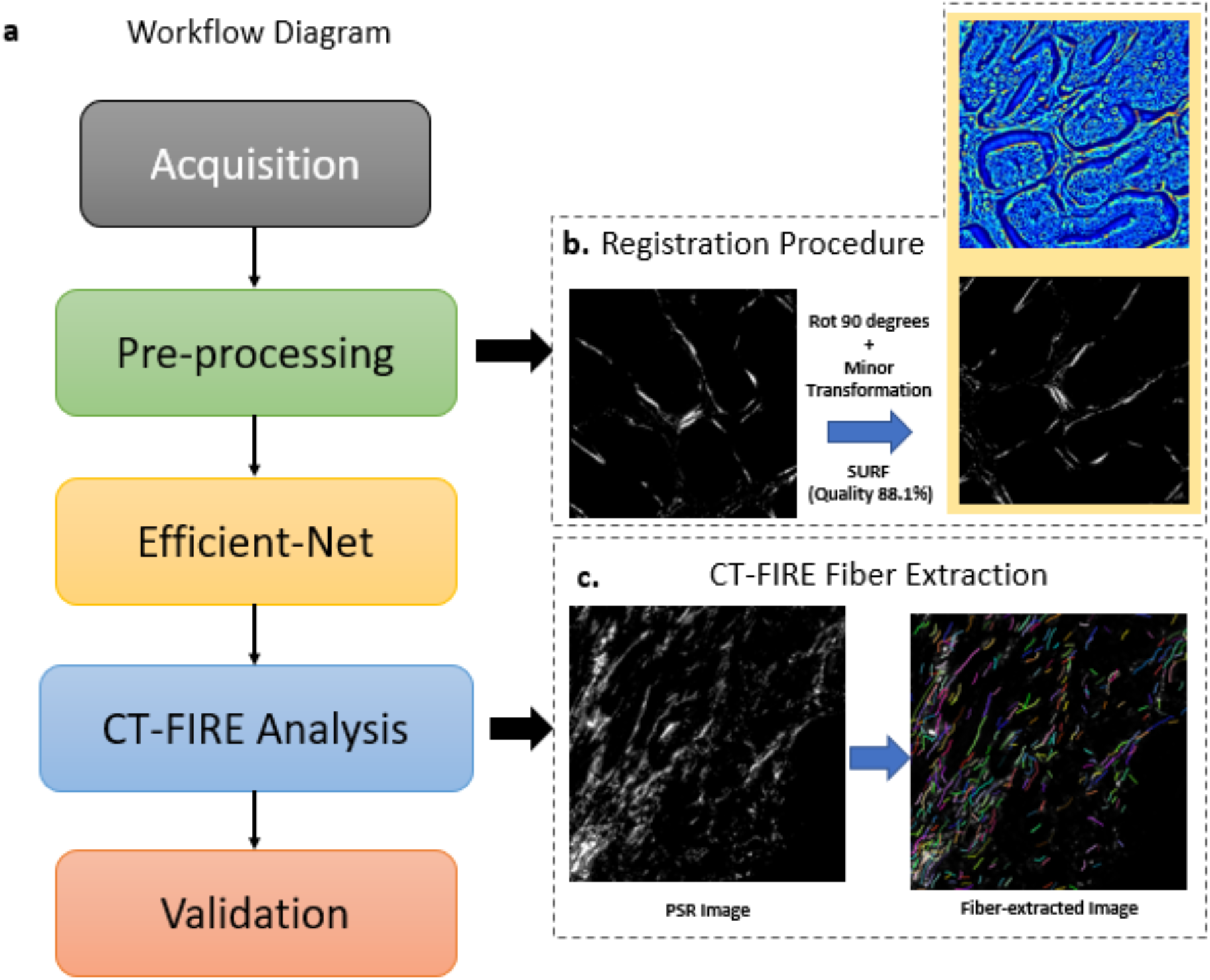
General workflow diagram of PICS training and validation. **a.** The standard procedure for PICS implementation begins with acquiring the SLIM and PLM images. Upon pre-processing, the image pairs are used to train the E-U-Net. Lastly, the inference from the E-U-Net is further analyzed using CT-FIRE and fiber statistics are analyzed as part of the validation process. **b.** Following manual 90 degrees rotation of PLM images, we use the MATLAB registration algorithm (SURF algorithm, specifically) to finely obtain the transformation matrix to be applied over the entire PLM images. **c.** CT-Fire extraction process is done using MATLAB with adjustable intensity and length thresholding strategy appropriate for each ground truth and prediction image dataset.

### Efficient U-Net

In order to detect solely the collagen fibers from the SLIM image, we trained a convolutional neural network with SLIM as input and PLM images as the ground truth. We used a modified U-Net, referred to as the Efficient U-Net (E-U-Net) **[67]**. The encoding path of E-U-Net is the EfficientNet in which the network was pretrained on image dataset from ImageNet **[73]**. Such modification reduces the complexity of the original U-Net,, while maintaining high accuracy **[74, 75]**. The detailed depiction of the overall network architecture of E-U-Net and the convolutional layer diagram of the EfficientNet encoding path are illustrated in **Figures 4a and 4b**, respectively. The learning rate is shown in Fig. 4c.

**Figure 4:**
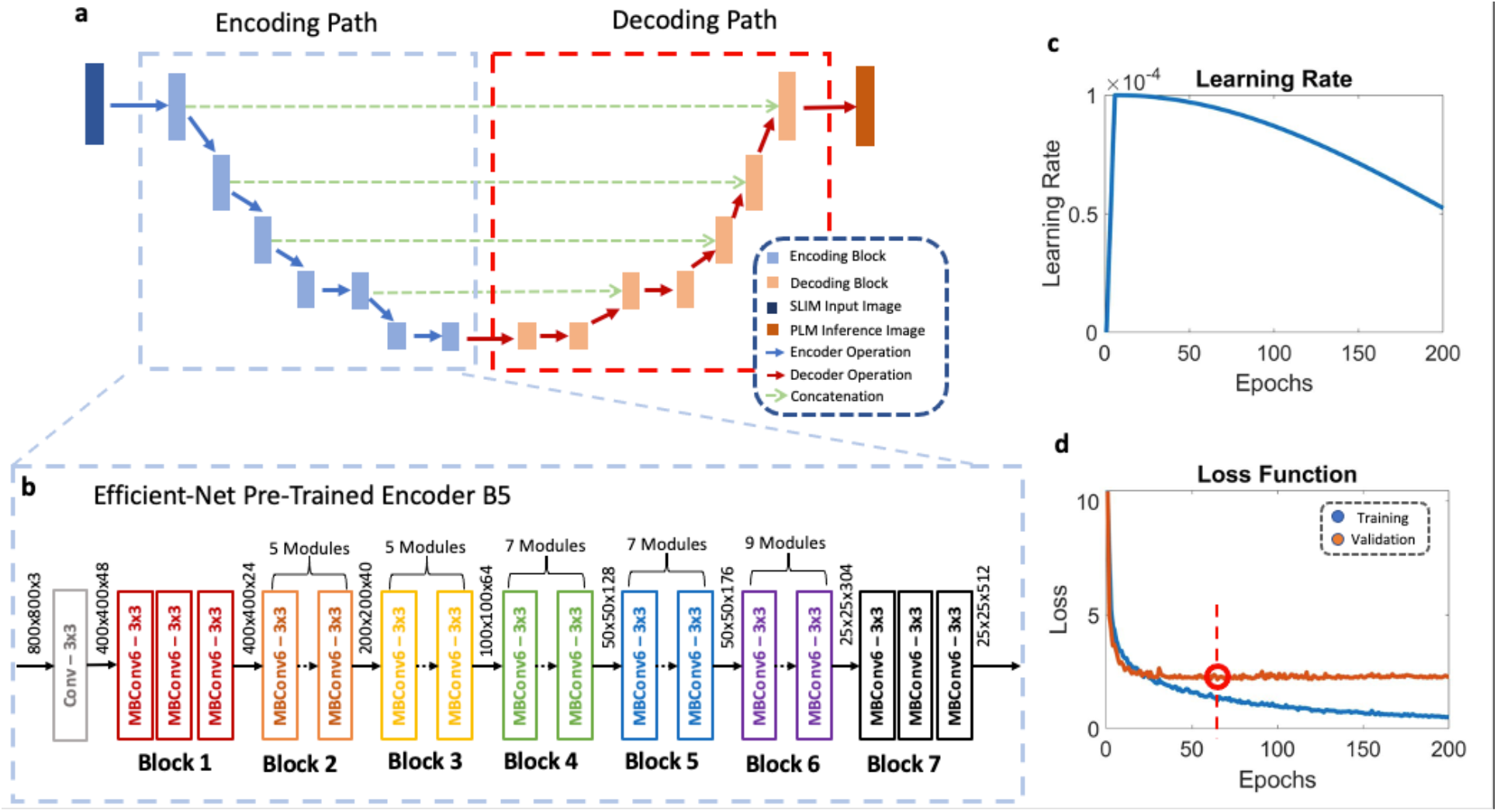
The network description. **a.** Schematic representation of the encoding and decoding path. The encoding path involves a pretrained Efficient-Net encoder. **b.** Detailed block structure of Efficient-Net B5 used for training. The encoder consists of one regular convolution layer (labeled in gray) and seven distinct blocks consisting of inverted mobile bottleneck (MB) convolution operations. **c.** Plot of the cosine learning rate over training epochs. **d.** Training and validation loss curve over epochs trained. The best validation loss was attained at epoch 65 (Labeled accordingly).

From the 710 acquired SLIM-PLM pairs, we trained the E-U-Net network with 496 SLIM-PLM pairs of images and used 142 pairs as validation set. We used a linear combination of two different loss functions, namely, the mean absolute error loss and Pearson correlation loss to find the updated model weights that best optimize the loss function. We used the EfficientNet B5 network as the backbone encoder of our E-U-Net and ran it for 200 epochs. The minimum value of the loss function was attained at epoch 65 as shown in **Figure 4d**, which we used to evaluate the accuracy metrics over the validation and test data sets. We discuss in more detail the training parameters and the interpretation of the network structure in the Methods section.

### Evaluation of Trained Model over the Test Set

To assess the inference performance, we applied the best model described above over 72 test datasets containing SLIM and PLM images. These 72 pairs were kept separate from the training set and, thus, were not “seen” by the E-U-Net. **Figure 5a** illustrates three SLIM core images. **Figures 5b and 5c** show the ground truth and PICS prediction, respectively. Using the 142 validation images to assess the network performance, the training script evaluated the average peak signal-to-noise ratio (PSNR) and structural similarity index measure (SSIM). The PSNR and SSIM values for the PICS inference were 31.7 and 0.75, respectively. The SSIM value from our model is within the acceptable level of performance according to previous reports **[76]**. One of the key features of both the original PLM image and PICS inference is that the glandular regions disappear, since glands are regions rich in epithelial cells that do not exhibit birefringence.

**Figure 5:**
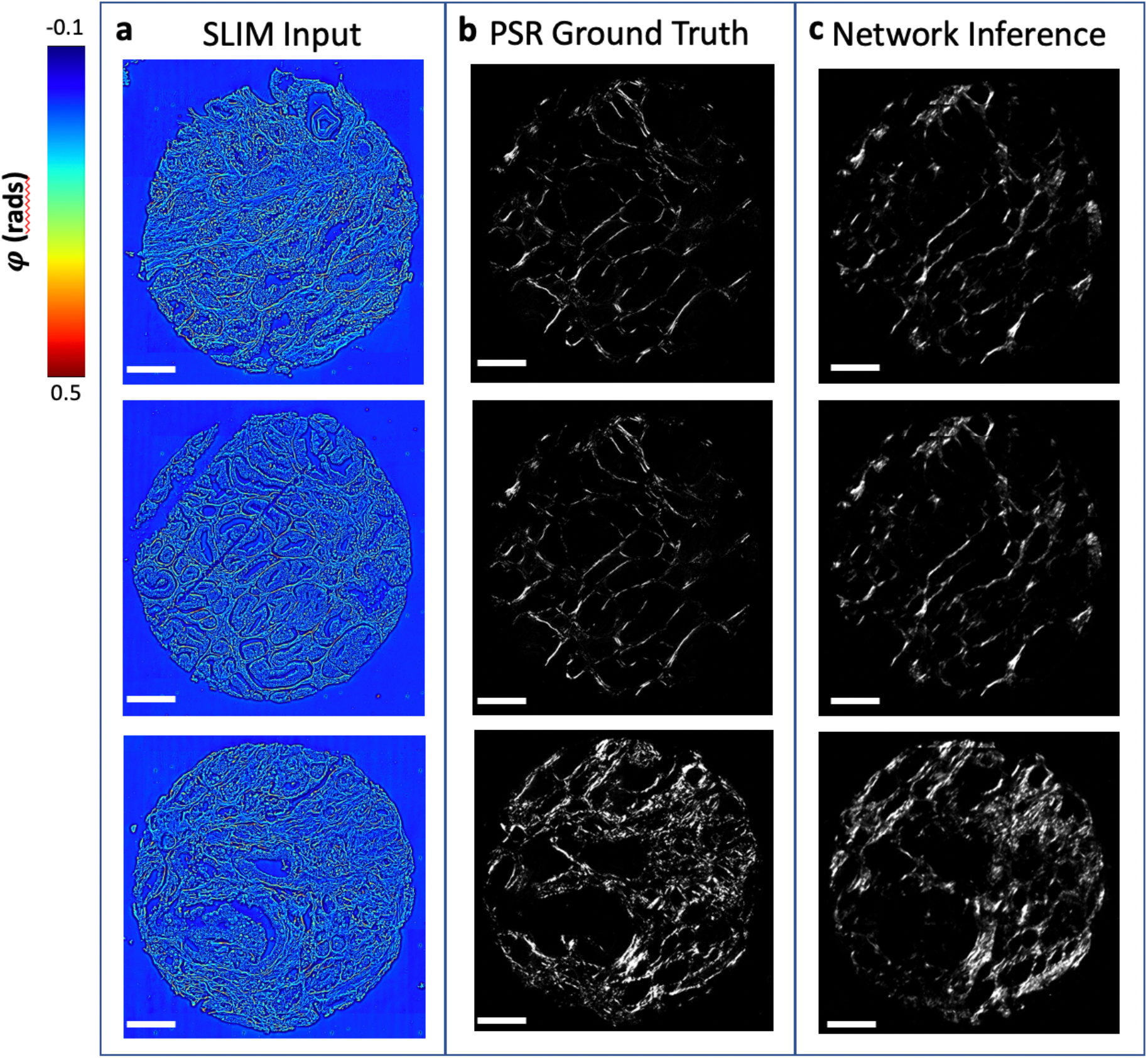
Result of E-U-Net performance over test dataset. **a.** SLIM input image of cores input to the network. **b.** The corresponding PSR ground truth images. **c.** PICS prediction of PSR signal. The white scale bar for each image represents 200 microns in length.

### CT-FIRE Results on Test Datasets

In order to validate PICS for the stain-free orientation analysis of collagen, we used the CT-FIRE (Curvelet Transform – Fiber Extraction) algorithm, originally used to localize individual collagen fibers from second harmonic generation microscopy images **[19].** Previous applications indicated that the CT-FIRE algorithm can also be applied to PLM images **[69]** and even SLIM images **[68]**, assuming that the data contain isolated collagen fibers. CT-FIRE takes advantage of the curvelet transform algorithm to create a segmented map consisting of detected collagen fibers from a given input image (**Figure 3c)**. We extracted individual collagen fibers from each of the test ground truth and the PICS inference and evaluated the statistics of the angular orientation, straightness, and the length associated with each localized fiber, as shown in **Figures 6a** and **6b**. In **Figure 6b**, we plotted the histogram distributions for fiber angle, length, and straightness associated with both the ground truth PLM and PICS output, as indicated. The histogram comparison shows strong overlap, indicating that PICS information can be used for analyzing collagen fibers directly on unlabeled specimens.

**Figure 6:**
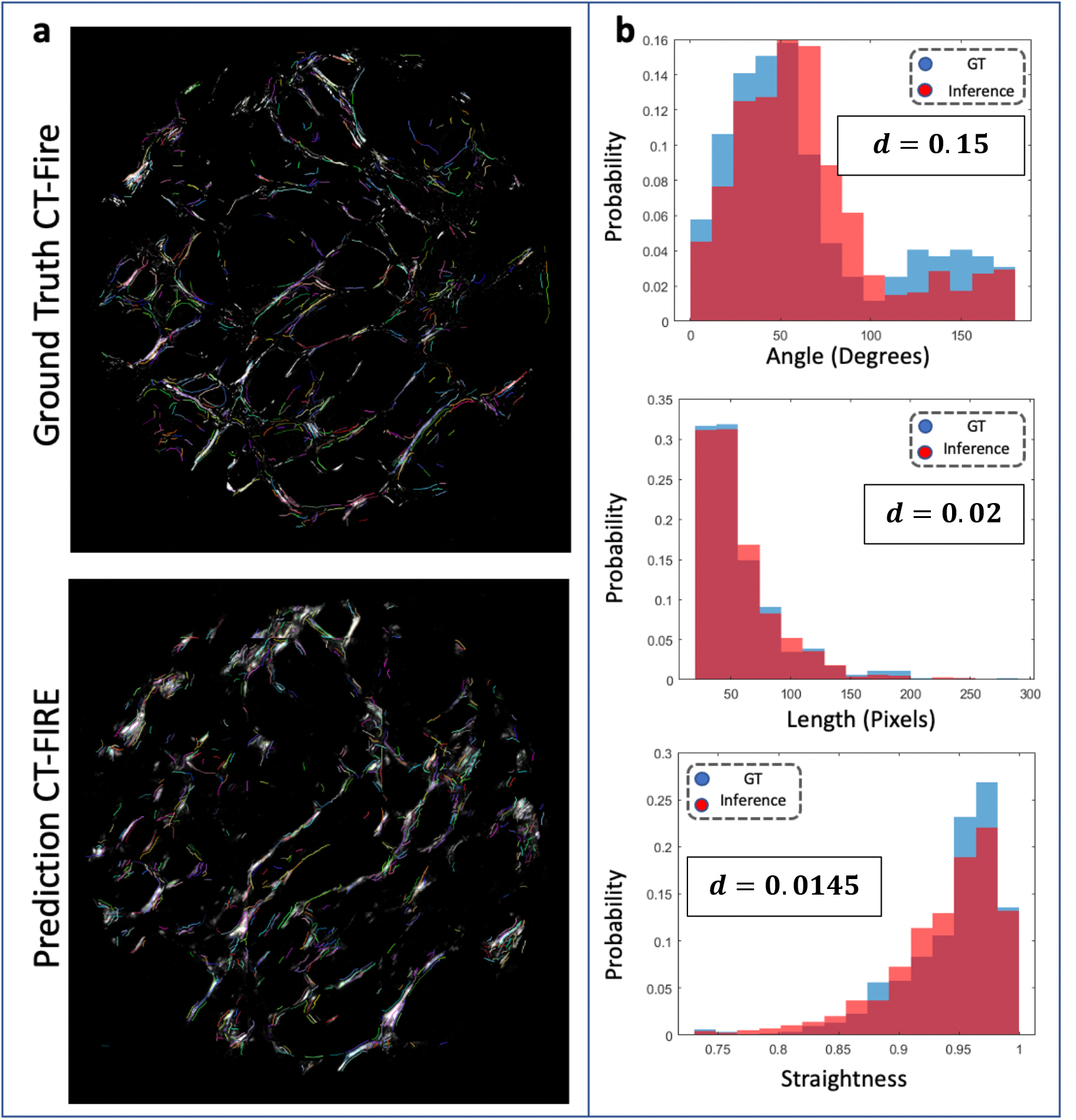
Result from the CT-FIRE analysis. **a.** Mapped collagen fibers on ground truth and prediction images. **b.** Discrete probability distribution for length, angle, and straightness of detected collagen fibers. The significantly small KL (Kullback-Leibler) divergence denoted in the bar graphs indicate statistical similarity of two red and blue bars graphs.

### Validation of CT-FIRE Statistics

In order to validate our CT-FIRE results, we applied a two-distribution t-test at 5 percent significance level over each of the 72 pairs consisting of ground truth and the corresponding PICS output for length, angle, and straightness (**Fig. 7b).** The KL (Kullback-Leibler) divergence was used to evaluate statistical similarity between the histograms for ground truth and PICS inference. From this quantitative result, we can conclude that PICS inference of the PLM image from SLIM input can be used for clinical inference in the absence of a physical stain.

## Methods

### Tissue Microarray (TMA)

The tissue microarray (TMA) used in this study was obtained from 217 prostate cancer patients, with four 0.6 mm cores sampled per prostatectomy specimen. Gleason grading was performed by pathologists on 192 cases using the 2019 International Society of Urological Pathology (ISUP) modification **[77]**. Of the 192 cases analyzed, 67 had modified Gleason score of 5-6, 103 scored 7, and 22 scored 8 to 10. Each tissue sample was deparaffinized prior to the staining process. The deparaffinized cores were then stained with picrosirius red and H&E and cover-slipped at the University of Illinois at Chicago.

### SLIM System

SLIM imaging is achieved by connecting an external SLIM module (CellVista SLIM Pro, Phi Optics, Inc.) on the side port of a phase contrast microscope **[49]**. The SLIM principle of operation is presented Ref. **[25]** and recently reviewed in Ref. **[49]**. In short, the SLIM module consists of a 4f lens system **[78]** and a space light modulator (SLM) in its Fourier plane to apply *π*/2, *π*, 3*π*/2, 2*π* phase shifts in addition to the existent *π*/2 phase shift applied by the phase contrast objective ring. From a set of four phase contrast images, we compute the quantitative phase delay from the sample. For SLIM, we used the 40x phase contrast objective with 0.75 numerical aperture and collected the images using an ORCA-Flash 4.0 CMOS camera (Hamamatsu). We set both the exposure time and the SLM stabilization time at 30ms.

### PLM System

PLM image setup is shown on the right port of the microscope in Figure 1a, which is essentially brightfield microscopy with the sample between two orthogonal polarizers. We placed the first linear polarizer on top of the condenser module of the inverted microscope and the corresponding analyzer right before the camera. Thus, the light from isotropic regions of the tissue undergoes full extinction upon encountering the cross-polarized analyzer. The PLM only transmits the light emerging from the birefringent regions of the tissue sample, mainly the PSR-stained collagen. For the PLM acquisition we used a 40x/0.75NA bright-field objective, which is consistent with the phase contrast objective used in SLIM. The PLM image was captured using a Neo CMOS camera (Andor) with 100 ms acquisition time.

### Image Registration

Image registration is very important for our image-to-image transformation using the convolutional neural network. Differences in the fields of view between the input SLIM image and the ground truth PLM image may arise from slight shifts and rotational misalignments between the two cameras in Figure 1. We used the MATLAB image registration estimator to correct such geometric misalignment. The estimator takes in two images, one as a fixed reference and the other as the image to be transformed to match the prior. Then the estimator outputs the transformation matrix which incorporates all types of transformations such as rotation, scaling, and affine. MATLAB image registration estimator offers various algorithms for feature recognition between the pair, as well as tunable parameters for optimization processes.

In order to maximize the performance of the feature recognition involved, we used two phase-contrast microscope images captured by both the left and right cameras. We assigned the image from the SLIM camera (on the left side of Figure 1) as the fixed reference image and the corresponding image from the PLM camera (on the right side of Figure 1) as the image to be transformed. The SURF (Speeded Up Robust Features) algorithm was selected for feature detection algorithm. The SURF algorithm detects significant features using the determinant of the Hessian of the input pair **[79]**. Then, the detected features are juxtaposed by the estimator to approximate the transformation matrix. Assuming mechanical stability, we applied the calibrated transformation matrix on every acquired PLM image prior to the network training.

### Efficient-UNet Network and Training

Efficient-UNet (E-U-Net) is a convolutional neural network originated from the regular U-Net **[80]** that infers the PLM counterparts of the input SLIM images without the presence of histological staining. The main difference between the U-Net and the E-U-Net is that E-U-Net uses a more efficient pre-trained encoder, EfficientNet, in the encoding path **[67]**. EfficientNet is a compact convolutional neural network that retains its very high performance while lowering the number of total network parameters being used in order to increase the convergence rate. Branching off from the EfficientNet B0, the least complex, more complex architectures were built ranging from B1 to B7 by adding more layers for higher performance **[67]**. We empirically selected the best performing EfficientNet architecture, EfficientNet B5, as the pre-encoder of our E-U-Net training session by juxtaposing the minimum loss function value attained by each architecture.

The dimensions of the images used to train the network were of 2048 by 2048, which is equivalent to 300×300 μm^2^ field of view. In order to reduce storage burden on the GPU hardware during training, we randomly selected 800 by 800 crops for every iteration of the training session. The model was trained with a total of 37,468,673 parameters and the ADAM optimizer to minimize loss functions **[81]**. We utilized a linear combination of mean absolute error loss and Pearson correlation loss function to assess both the training loss and the validation loss presented in Figure 5b. We used the mean absolute error (L1) over the mean squared error (L2) since Mean Absolute Error (MAE) promotes strict boundaries, while MSE allows more loose cutoff boundaries **[82]**. Having a strict boundary is very important especially when we need to extract morphometric information per collagen fiber in the CT-FIRE analysis. The learning rate was designed such that the rate of convergence decreases with a diminishing cosine profile **[83]** as shown in **Figure 5c**, starting from the initial value of 10^−4^. We trained the network for a total of 200 epochs and selected the neural network model that yielded the least loss function labeled with a red circle in **Figure 5b**.

We used the NVIDIA GeForce RTX 2080 GPU hardware throughout the E-U-Net training under CUDA 10.1. All scripts used for training and validation were written and run using Python 3.6 and TensorFlow 1.14.0.

### CT-FIRE Analysis

We used the CT-FIRE algorithm, as described by Bredfeldt et al. **[19]**, to automatically extract the fibers for the ground truth and corresponding PICS inference result for each field of view. The algorithm of CT-FIRE first starts by performing a 2D curvelet transform on the images, which creates a visualization of both the position and the spatial frequency **[84, 85]**. The main use of curvelet transform is to emphasize the line-like features and suppress background noise within a given image **[19]**. The algorithm further performs a transform to show the distance from a given pixel to the background. By tracing the path of maximum distance, the algorithm extracts list of line-like features including what is referred in the original CT-FIRE literature as the nucleation points **[19]**. Lastly, the algorithm removes some fibers that may be either too short or too wide to present a colored segmentation map of detected fiber strands (see **Figure 3c)**. We use three main parameters (length, straightness, and angle) associated with all fibers detected from the algorithm. Our results indicate significant similarity between the CT-FIRE output generated on the ground truth and PICS images, as shown by performing the two-distribution t-test for each metric.

For the validation process, we ran the CT-FIRE script over 72 pairs of ground truths and corresponding PICS predictions unseen by the trained network to run the two-distribution t-test over length, angle, and straightness separately. The null hypothesis used in the t-test is that the two distributions are like the ones between the ground truth and prediction. We used 5% (0.05) for the statistical significance threshold.

## Summary and Discussion

We showed that PICS allows label-free, non-destructive collagen imaging and characterization, without the need for picrosirius red stain or SHGM imaging. Central to this accomplishment is SLIM’s capability to yield a quantitative assessment of tissue morphology with high sensitivity. We used the PICS method to infer the PSR-stained image from the SLIM dataset as input. The PSNR and SSIM evaluated over the unseen test data set were 31.7 and 0.75, respectively. Upon network inference, we used the CT-FIRE algorithm to extract morphological statistics on the individually identified fibers.

While SLIM measurement yields label-free quantitative phase images of the complete tissue architecture, PLM, with the aid of picrosirius-red stain, provides images specifically of collagen fibers. PICS combines the advantages of the two modalities through the power of deep learning. Label-free imaging of collagen with high specificity can be performed without staining or other preparation beyond tissue slicing. Since collagen imaging and analysis has a variety of applications ranging from cancer pathology **[3, 4, 6, 19, 86, 87]** to skin aging **[88, 89]** and wound healing [90], we anticipate that PICS inference of collagen fibers can be implemented on unstained tissue samples from various organs. Since SLIM can be implemented as an upgrade to an existing phase contrast microscope, and the deep-learning algorithms can run on a typical PC, it is likely that this approach can be adopted broadly in clinical practice.

